# Identification of novel circadian transcripts in the zebrafish retina

**DOI:** 10.1101/416222

**Authors:** Soundhar Ramasamy, Surbhi Sharma, Bharat Ravi Iyengar, Shamsudheen Karuthedath Vellarikkal, Sridhar Sivasubbu, Souvik Maiti, Beena Pillai

## Abstract

High fecundity, transparent embryos for monitoring the rapid development of organs and the availability of a well-annotated genome has made zebrafish a model organism of choice for developmental biology and neurobiology. This vertebrate model, a favourite in chronobiology studies, shows striking circadian rhythmicity in behaviour. Here, we identify novel genes in the zebrafish genome, which shows their expression in the zebrafish retina. We further resolve the expression pattern over time and assign specific novel transcripts to the retinal cell type, predominantly in the inner nuclear layer. Using chemical ablation and free run experiments we segregate the transcripts that are rhythmic when entrained by light from those that show sustained oscillations in the absence of external cues. The transcripts reported here with rigorous annotation and specific functions in circadian biology provide the groundwork for functional characterisation of novel players in the zebrafish retinal clock.

## Introduction

Circadian rhythms are physiological and behavioural changes that recur in about 24 hours that allows the organism to adapt to fluctuations in the external environment driven by the day and night cycles imposed by the sun. Empirical observations in natural conditions and elegant experiments in laboratory models established the three tenets of circadian biology: (a) the core clock is internal (b) it can be entrained to external cues, especially light (c) the basic clock mechanism is conserved. The internal nature of the clock became evident when it was seen that the circadian expression of certain genes and the manifestation of circadian behaviour like sleep or eclosion would continue to show approximately 24 hour rhythms irrespective of the external cues. Thus, under constant dark conditions imposed externally, the fruitfly, *Drosophila melanogaster*, will continue to display nearly 24 hour rhythms, gradually and inexorably slipping from the objective time. However, in the presence of external light cues, the rhythm adjusts to match the cue(Partch et al. 2014).

Transplantation experiments in model organisms (Ralph et al.1990) and the prevalence of free-run circadian rhythms in certain types of blindness(Sack et al.1992) have now firmly established that the core clock resides in the suprachiasmatic nucleus (SCN) in the mammalian brain. Many organs and even individual cells have peripheral clocks that are synced to the master clock but when isolated, continue their individual, but eventually dampening, rhythms(Tosini and Menaker 1996; Moore and Whitmore 2014). In organisms as distant as the fungus *Neurospora crassa*, insects like the fruitfly, nocturnal rodents like mice, and humans, the primary clock mechanism depends on feedback regulation of a set of highly conserved genes(Loudon et al. 2000). The various manifestations of circadian rhythms are brought about by the expression of output genes, under the regulation of the core regulatory transcription factors. The core clock genes and its associated transcriptomes are highly organ-specific (Zhang et al. 2014).

The retina is a thin layer of complex tissue along the inner wall of the eye, made up of intricately connected layers of cells with more than 60 functionally distinct neuronal types (Masland 2012). The rods and cones, with their distinctive shapes, collect the input light and communicate these signals to bipolar cells in the inner nuclear layer (INL). From INL, the signal is transmitted to the retinal ganglion cells (RGCs) in the ganglion cell layer (GCL) which eventually relay it to the brain via the optic nerve. Further, signals are modulated by amacrine and horizontal cells when its flows from photoreceptors to bipolar and bipolar to RGCs respectively. Since the retina is an extension of the central nervous system many phenomena relevant to CNS disorders have been studied in the more accessible retinal cells (Athanasiou et al. 2013; Lamb et al. 2007). Retinal cell types are prone to oxidative damage and endoplasmic reticulum (ER) stress arising from the rapid turnover of proteins besides, light-induced damage to lipids, protein and DNA. This necessitates extensive repair processes and renewal in coordination with circadian fluctuations. Though these repair processes are shared across evolution, the zebrafish retina is unique in its ability to regenerate all cell types upon extensive damage even into adulthood, an ability attributed to retinal Muller glial cells (Jorstad et al. 2017).

The Muller glial mediated injury response is marked by the strong up regulation of core clock genes in zebrafish but not in mouse (Sifuentes et al. 2016). Besides this, many vital functions of the retina are under the influence of diurnal or circadian factors; melatonin production (Huang et al. 2013), photoreceptor disc shedding(Grace et al. 1999), extracellular pH of the eye (Dmitriev and Mangel 2000) and rod-cone coupling (Ribelayga et al. 2008) to name a few. The segregation of genuinely circadian genes from those that merely respond to light and hormonal cues will significantly enhance our knowledge of circadian biology and the role of the retina in providing inputs to the core clock and maintaining a robust peripheral clock. Further, an extensive catalogue of temporally resolved retinal gene expression in zebrafish would be a valuable resource for exploring the gene regulatory networks behind the complex multicellular organisation of retinal function.

We compared the expression profiles of core clock genes in zebrafish, mouse and baboon to establish that zebrafish recapitulates the salient features of the diurnal behaviour of the baboon. Since temporal gene expression patterns of the zebrafish retinal tissue were not available, we generated RNA-seq data and carried out extensive annotation of the retinal transcriptome of the zebrafish, during the light and dark phases of a day. Further, we validated the expression of selected novel transcripts and studied their response to light and dopamine. Lastly, we used free run experiments under constant dark conditions, to unambiguously assign a circadian role to a subset of the novel transcripts. This study not only provides a valuable resource for understanding the zebrafish retina but also throws up novel genes, that code for proteins and non-coding RNAs whose functional role in circadian gene expression is currently unknown.

## Material and methods

### Zebrafish maintenance

Assam wild type (ASWT) strain was used for all the above experiments. Adult fish used were six months to one year old. Maintenance and experiments were strictly within the guidelines provided by Institutional Animal Ethics Committee (IAEC) of the CSIR Institute of Genomics and Integrative Biology, India with timed feeding and uniform temperature at 28°C±2°C. ‘The Zebrafish Book-a guide for the laboratory use of zebrafish’ was used for recommended standard practice (Westerfield). Zebrafish were anaesthetized using Tricaine (Sigma, USA) for whole eye cup dissection. Dim red light was employed for eye cup dissection at dark time points.

### Library preparation and RNA-seq data analysis

From zebrafish whole eye, RNA was isolated using RNeasy kit (Qiagen).Approximately, 1µg of total RNA was used for library preparation using Truseq stranded RNA library. (Illumina Inc. USA). Ribosomal RNA (rRNA) in each sample was depleted as per manufacturer’s instructions (Illumina Inc. USA) followed by fragmentation. Complementary DNAs (cDNAs) were prepared after rRNA depletion. Further, sequencing adapters were ligated and the enriched libraries were subjected to 101x 2 Paired End sequencing using Hiseq 2500 platform following regular protocol (Illumina Inc. USA). The FASTQ sequencing reads were adapter-trimmed along with a minimum length cut-off of 50 bases using prinseq-lite. The reads were aligned to zebrafish genome assembly (Zv9) using TopHat (v2.0.11) followed by reference based assembly by Cufflinks (v2.2.1). Then differentially expressing transcripts were identified using Cuffdiff (v2.2.1).

### qRT-PCR analysis

Tissue or whole animal RNA was isolated using RNeasy kit (Qiagen). cDNA was prepared using random nanomer based reverse transcriptase core kit (Eurogentec).qRT-PCR was carried out using SYBR Green master mix (Applied Biosystems) for detection in Light cycler LC 480 (Roche). All primers used for qRT-PCR are given in table.

### Th+ amacrine cell chemical ablation

Depletion of Th+ amacrine cells in one eye was achieved by 2μl intraocular injection of mixture of 1:1 6-hydroxydopamine (6-OHDA) and pargyline (Sigma) for two consecutive days at ZT4. Corneal incision and intra-ocular injection were followed as described in (Li and Dowling 2000). After one week of injection, Th+ amacrine cells ablation in retina was analysed by immunolabeling and qRT-PCR against tyrosine hydroxylase protein and mRNA respectively.

### Whole retinal flat-mount immunolabeling

Whole retina was dissected out of the eye cup; specimens were fixed in 4% paraformaldehyde and 5% sucrose for overnight followed by 0.1M glycine treatment for 10min at room temperature. For antigen retrieval, tissue was placed in sodium citrate buffer (10mM sodium citrate, 0.05% Tween-20; pH-6) and incubated in boiling water for 10 minutes and cooled to room temperature followed by washing with 0.5% Triton X 100 in PBS, blocking for 2 hrs (5% BSA,1% DMSO, 1% Tween, 0.1% Sodium azide in PBS), primary antibody (1:500,tyrosine hydroxylase, Sigma Aldrich, SAB2701683) in blocking buffer with 2% BSA was incubated for overnight at 4^°^C and secondary antibody (1:500, FITC conjugated, Invitrogen) for 2hrs at room temperature. Tissue was mounted and imaged using Confocal Leica TCS SP8 Confocal microscope.

### RNA *in situ* hybridization

WISH was performed by following the standard zebrafish protocol (Thisse and Thisse 2008) While, RNA in situ on adult eye cryosection were subjected to an extra step of acetylation treatment; slides with sections were treated with 0.25% acidic anhydride in 0.1M ethanolamine (pH-8) to prevent non-specific binding of probe on slides.

### Single cell transcriptome analysis

Publically available dataset GSE106121 was used in our analysis. Custom code provided by authors (https://bitbucket.org/Bastiaanspanjaard/linnaeus.) was used in single cell transcriptome analysis using Seurat (Macosko et al. 2015; Butler et al. 2018)

## Results

Circadian oscillations in gene expression have been documented in different tissues, sub-regions of the brain (Zhang et al. 2014; Mure et al. 2018)in a variety of organisms (Boyle et al. 2017; Kuintzle et al. 2017; Hatori et al. 2014). We compared the reported gene expression profile of 5dpf (days post fertilization) old zebrafish embryo (Li et al. 2013) and liver of the adult fish (Boyle et al. 2017) to recently reported data from baboon (Mure et al. 2018) and mice (Zhang et al. 2014). We found that many key circadian genes showed similar peak time in zebrafish and baboon, but not in the nocturnal rodent model (Figure1A). Mure et al. showed overall transcriptional activity was muted in the baboon liver in the early night. In a close agreement, zebrafish liver also shows a biphasic pattern of transcriptional activity (Figure1B). Besides these similarities to the circadian regulation of diurnal mammals, zebrafish offers additional advantages like a conserved retinal architecture, rapid development and large clutches of transparent embryos,. Despite the extensive work on the zebrafish retina (Gestri et al. 2012; Link and Collery 2015; Avanesov and Malicki 2010) and circadian behaviour (Cahill 2002; Hirayama et al. 2005), there is no catalogue of temporally resolved gene expression patterns in the zebrafish retina.

**Figure 1:**
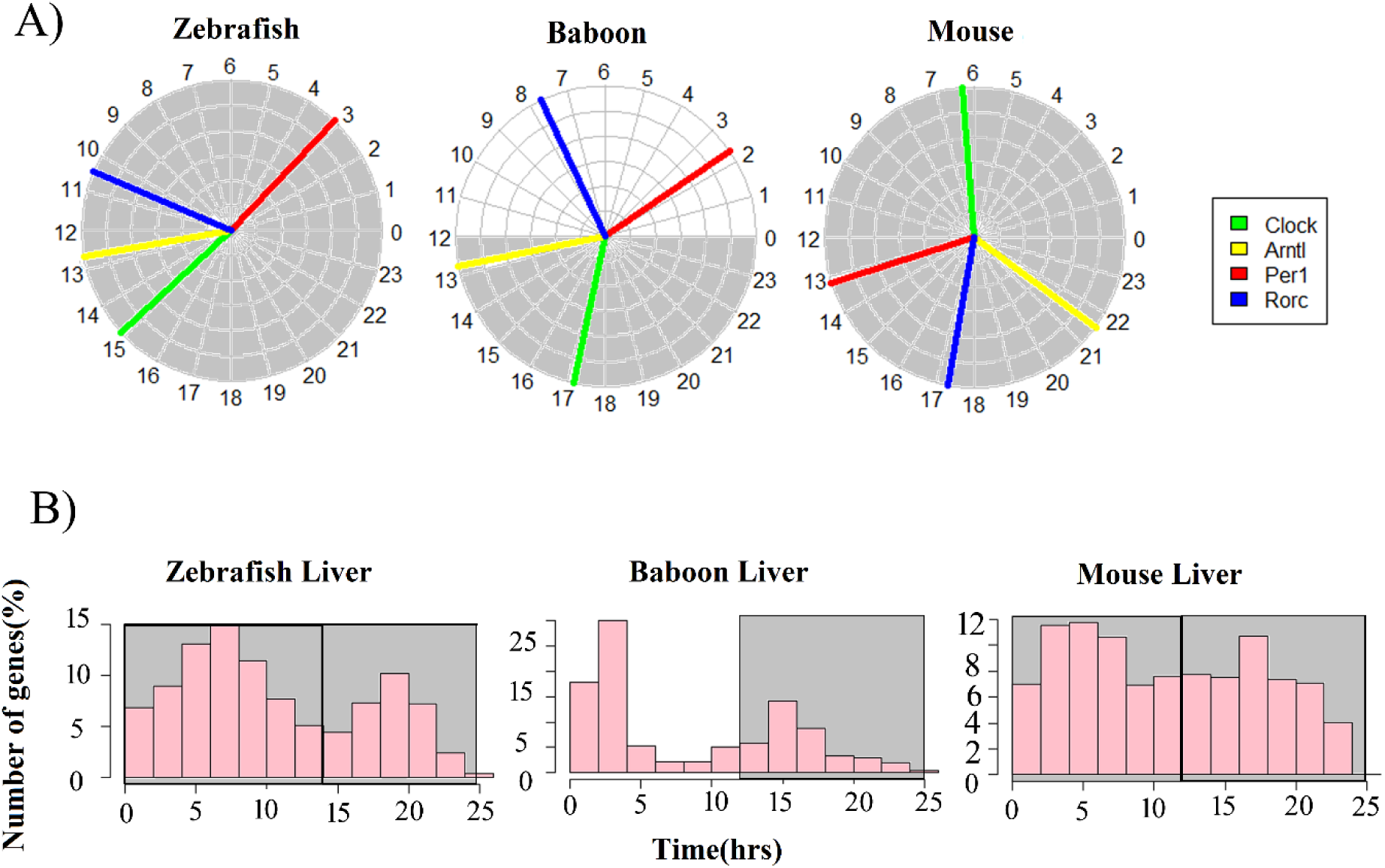
Zebrafish core clock gene expression resembles that of the diurnal baboon: (A) Peak expression phase of core clock genes in zebrafish (5dpf), baboon and mouse. Baboon and mouse datasets are average expression calculated from 64 and 12 different tissues respectively. The dark phase of the lighting condition is shaded gray. (B) Phase distribution of liver circadian transcriptome of zebrafish, baboon and mice

We collected whole eyecups from adult zebrafish exposed to alternating light-dark cycles. Total RNA isolated from the whole eyecup was collected at two-time points: ZT4 and ZT16 (Figure.2A); ZT0 representing switch from dark to light and ZT14 is representing the switch from light to dark condition. RNA-Seq libraries constructed from the pooled total RNA of four eyecups at each time point was subjected to Illumina short read sequencing. Zebrafish Reference guided transcriptome assembly was performed using TopHat (V2.0.11)-Cufflinks (v2.2.1)(Trapnell et al. 2012)and novel transcripts with no previous annotations were identified (Figure S1). Expression values were assigned to both novel transcripts and the transcripts already annotated in the zebrafish genome version (Zv9). Transcripts were found to be more or less uniformly distributed across the genome. At the extremes, chromosome 5 has the highest and chromosome 24 has the lowest density of the transcripts, respectively (Figure 2B). In total, 39% (ZT4) and 31% (ZT16) of known genes were identified in our study suggesting that the zebrafish eye expresses a substantial diversity of genes. This is in agreement with previous reports of mouse and human retinal gene expression profiles(Storch et al. 2007; Pinelli et al. 2016).

**Figure 2:**
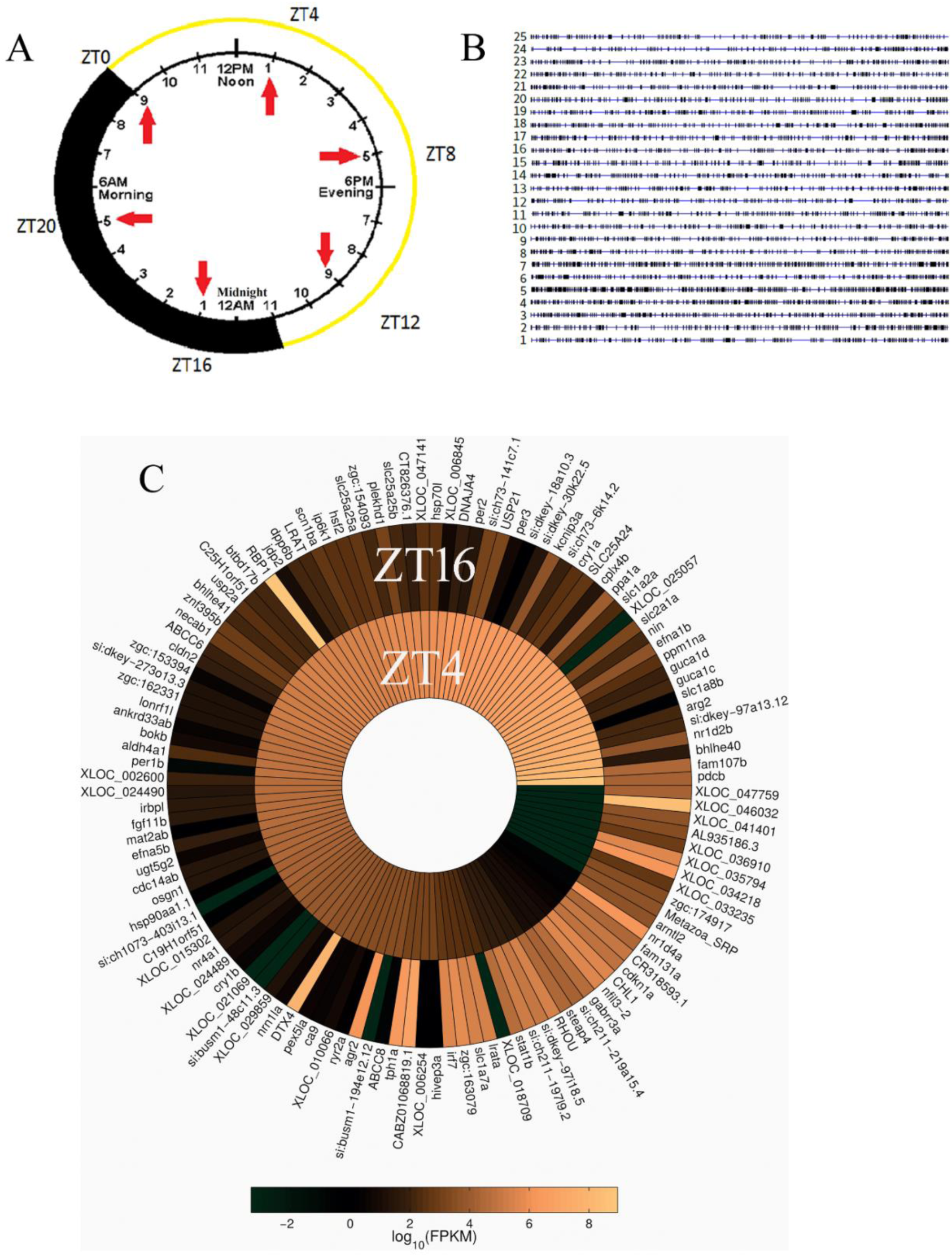
Diurnal transcriptome of adult zebrafish whole eye: (A) Schematic of sample collection time points and lighting duration (14 hrs light/10hrs dark) used in zebrafish maintenance. (B) Density of novel transcripts across the genome; each horizontal line depicts a chromosome and black boxes denote transcripts. (C) Heat map display differentially expressed transcripts from RNA-seq analyses of adult zebrafish whole eye from ZT4 and ZT16 time points

To segregate genes that show ultradian fluctuations, we compared the gene expression profiles of zebrafish eyes at ZT4 and ZT16, i.e. 4 hours after light (ZT4) and 2 hours after onset of darkness (ZT16). The transcripts with significant variation at the two time points were identified using Cuffdiff (v2.2.1). After imposing an arbitrary cut-off of 5 FPKM either at ZT4 or ZT16, 120 transcripts were found to be differentially expressed with 87 showing higher expression during light conditions, at ZT4; while the remaining 37 transcripts were induced during darkness, at ZT16 (Figure 2C). Ten transcripts amongst this set of diurnal eye transcripts are known to code for clock components (Table S1). Notably, per1b, per3, per2, cry1a, cry1b and nr1d2b showed higher expression during the day, in agreement with previous reports in zebrafish(Huang et al. 2018; Cahill 2002). Having established that the set of 120 differentially expressed genes show the expected diurnal regulation, we looked more closely at 21 novel transcripts. Seven novel transcripts showed low sequence complexity, making them unsuitable for further validation. The remaining 14 transcripts, named pc1 to pc14 were selected for validation by qRT-PCR. Despite repeated attempts with different primers, pc4 and pc9 could not be amplified consistently. Seven out of 12 tested transcripts (pc1, pc2, pc3, pc5, pc6, pc7& pc12) showed mutually corroborating differential expression in RNA-seq and qRT-PCR. However, two transcripts (pc13& pc14) showed discordance between qRT-PCR and RNA-seq (Figure 3A).

**Figure 3:**
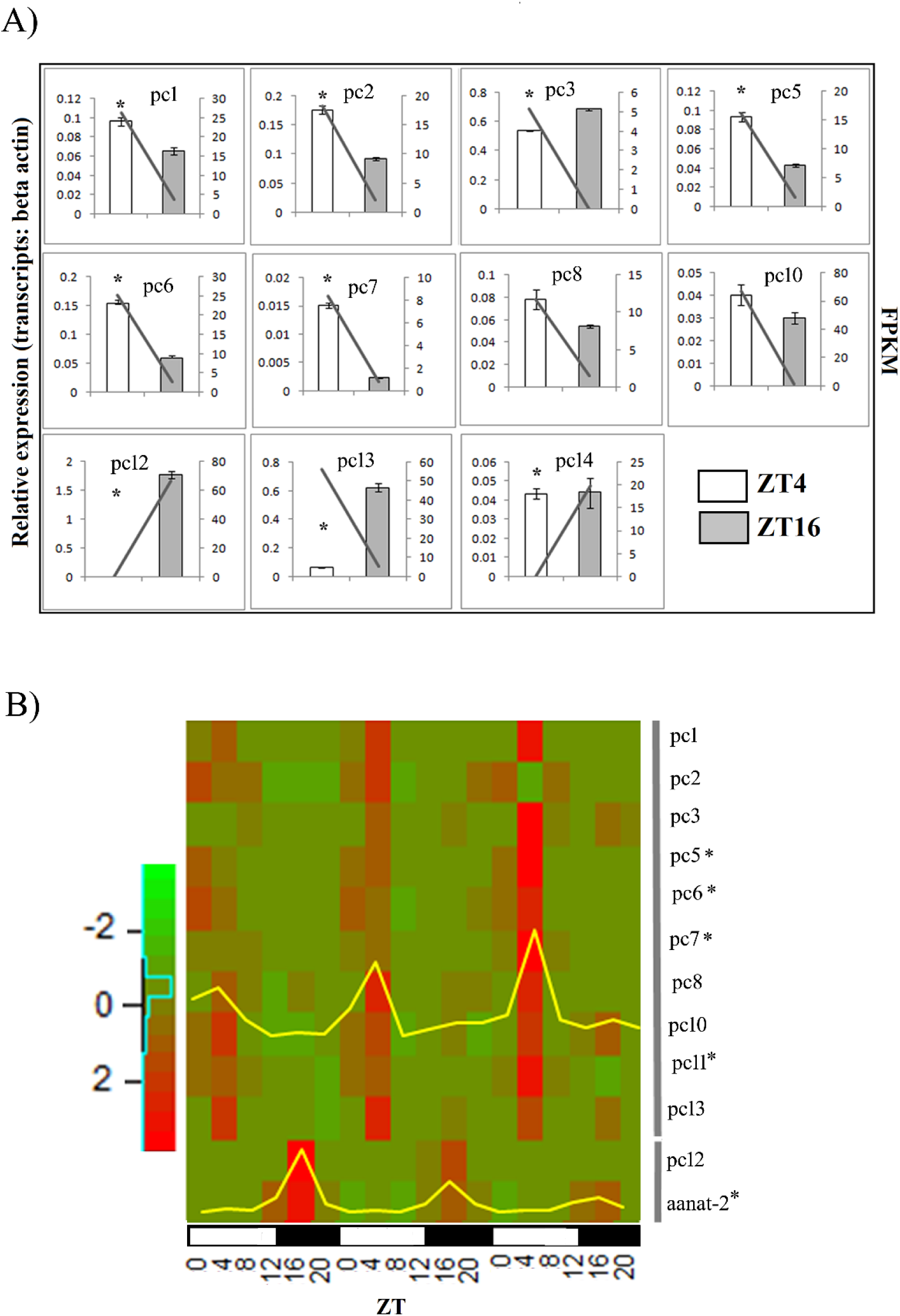
Differential expression of novel transcripts shows 24 hr diurnal rhythm : (A) qRT-PCR validation of novel transcripts RNA-seq differential expression trend. The values, normalised for beta-actin, are mean ± s.e.m. (n=5), with left and right y-axis representing qRT-PCR relative expression value and FPKM respectively. The line graph indicates the FPKM trend of ZT4 and ZT16. (B) Heat map displays temporally resolved novel transcripts expression from whole eye of adult zebrafish collected over 3 consecutive days. The values, normalised for beta-actin and scaled (n=1/time point). Yellow line overlay depicts average expression profile of the group of genes marked by grey vertical lines. * indicate the transcript rhythm statistical significance calculated using meta2d.

The RNA-Seq experiments and qRT-PCR experiments only sampled two time points, one each under light and dark conditions. To establish the diurnal rhythmicity of the selected transcripts, we repeated the qRT-PCR experiments using samples collected at four-hour intervals over three consecutive days with light and dark cycle entrainment (Refinetti et al. 2007). Most of the novel transcripts peaked at ZT4, although pc12, like the positive control gene arylalkylamine N-acetyltransferase-2 (aanat-2) (Gothilf et al. 1999; Falcón et al. 2001; Appelbaum et al. 2006), peaked at ZT16. This is in agreement with the pattern observed in the previous experiments since pc12 was strongly downregulated at ZT4 and upregulated at ZT16 (Figure 3B).

Next, we separated the retina comprising the thin inner layer of cells and in this fraction enriched for retinal cells; we studied the expression of the novel genes. All but pc14 showed enrichment in the retina. These genes showed a range of retinal enrichment, some (pc1, pc6 and pc8) even higher than that of the aanat-2 which served as a positive control (Figure 4A).

**Figure 4:**
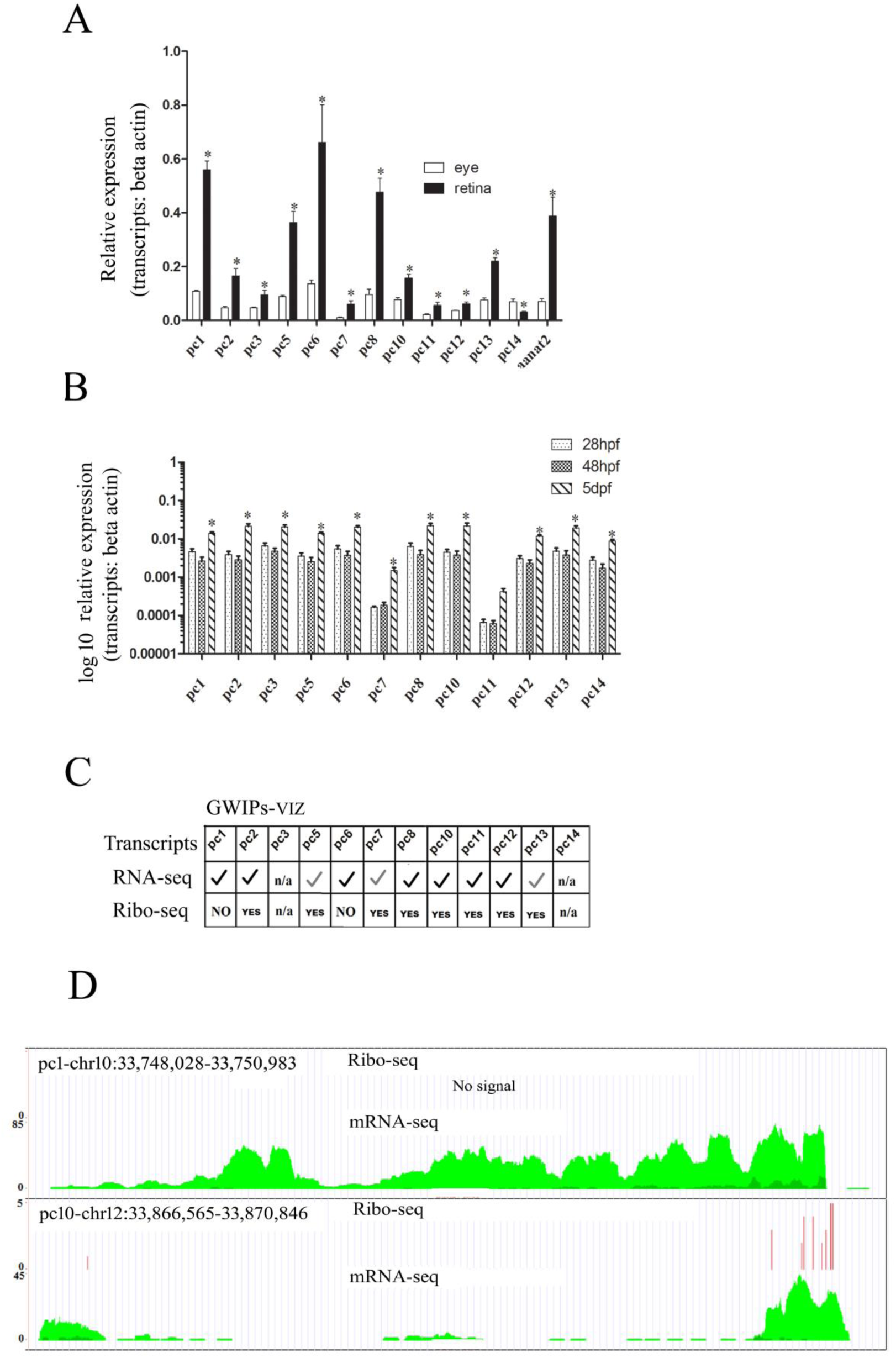
Ribo-seq status of retina enriched novel transcripts: (A) Novel transcripts enrichment between whole eye and retina of adult zebrafish isolated at ZT4. The values, normalised for beta-actin, are mean ± s.e.m. (n=5). (B) Relative expression (log10 transformed) of novel transcripts across early developmental stages - 28hpf, 48hpf and 5dpf of zebrafish. The values, normalised for beta-actin, are mean ± s.e.m. (n=5). * indicates the statistical significance (p-value<0.05) calculated between 48hpf and 5dpf. (C) The table display RNA-seq and Ribo-seq coverage from GWIPs-viz public datasets. Black tick (?) indicates the qRT-PCR trend coincides with RNA-seq coverage and grey tick (?) indicates discordance. (C) Ideogram of pc1 and pc10 transcripts indicating Ribo-seq coverage (red) and RNA-seq coverage (green). Ideogram was created using Gviz R packages.

In zebrafish, aanat-1 and aanat-2 are both expressed in the retina, but only aanat2 shows persistent circadian expression in constant darkness(Gothilf et al. 1999).

Next, we carried out qRT-PCR experiments at various developmental stages (28hpf, 48hpf, 5dpf) (Figure 4B) to identify the peak expression times of these transcripts during zebrafish retinal development. Behavioural responses to visual stimuli suggest that the zebrafish eye starts detecting light between 2.5 and 3.5 days post fertilization (dpf). It is well established that the zebrafish optic cups are clearly discernible in embryos 24 hours post fertilization (hpf) (Easter and Nicola 1996) while the period between 24hpf and 36hpf is marked by rapid cell division and the formation of post-mitotic neurons. By about 50hpf, the rod cells start expressing opsin genes and by 5dpf, all five types of photoreceptor cells: rods, short single cones, long single cones, and short and long members of double cone pairs are morphologically distinct. All the novel retinal transcripts, identified here, without exception, showed highest expression at 5dpf (Figure 4B), a stage at which, neurogenesis in the zebrafish retina is complete (Nawrocki et al. 1985).

With the rapid advances in transcriptome sequencing, it is now clear that protein-coding sequences account for only a small fraction of the genes in mammalian genomes. Although to a lesser extent, non-coding RNAs are also abundant in lower organisms where they carry out various functional roles in gene regulation. Ribo-seq profiling identifies the transcripts found in association with ribosomes, thus providing an experimentally verified resource of likely protein-coding transcript (Ingolia et al. 2011). We used publically available Ribo-seq data (Chew et al. 2013) to classify the novel transcripts identified in this study into potential non-coding RNA and protein-coding transcripts. The chromosome coordinates used to query the GWIPs-viz browser (Michel et al. 2014) that allows visualisation of the Ribo-seq data are listed in Table S4. The transcripts, pc1 and pc6 were detected in RNA-seq but not in the Ribo-seq suggesting that they are non-coding transcripts (Figure 4C, D). The RNA-seq / Ribo-seq data was inconclusive for pc3, pc4 and pc14, due to poor coverage in these experiments. Nine of the 14 transcripts (pc2, pc5, and pc7-13) were detected in Ribo-seq experiments, suggesting a protein-coding role. We translated the RNAs and identified predicted proteins arising from these transcripts. The largest novel protein, potentially coded from these transcripts was 1980 amino acid long and showing significant similarity to Ryanodine receptor2 (RYR2) at the protein level. In rodents, Ryr2is known to regulate circadian output from the SCN neuron(Aguilar-Roblero et al. 2007). These observations further reinforced that the newly identified transcripts play roles in circadian gene expression.

As mentioned before, one of the unique advantages of the zebrafish retina is the layered arrangement of cells, which facilitates the visualisation of cell types and spatial expression patterns by a combination of in situ hybridisation and nuclear staining in tissue sections. We carried out the whole mount in situ hybridisation (WISH) of the 5dpf embryo eye to detect the expression of the novel transcripts using labelled anti-sense riboprobes. Most of the transcripts, (pc1, 5, 6, 7, 8, 10, 13) showed clear expression pattern in the eye and brain, while no signal was observed with the use of respective sense probes (Figure 5A). Closer examination of the expression patterns in the retinal layers using cryo-sections showed that all but pc13 showed inner nuclear layer enrichment. pc13, a potentially protein-coding transcript was alone expressed in the retinal ganglion layer (Figure 5A). WISH on 5dpf embryos, and adult eye cryosections also revealed the diurnal rhythm in the expression of transcripts pc1 and pc10. Both these transcripts showed a rapid clearance of the transcript at ZT16 (dark) following high expression in the INL at ZT4 (light) (Figure 5B, C).

**Figure 5:**
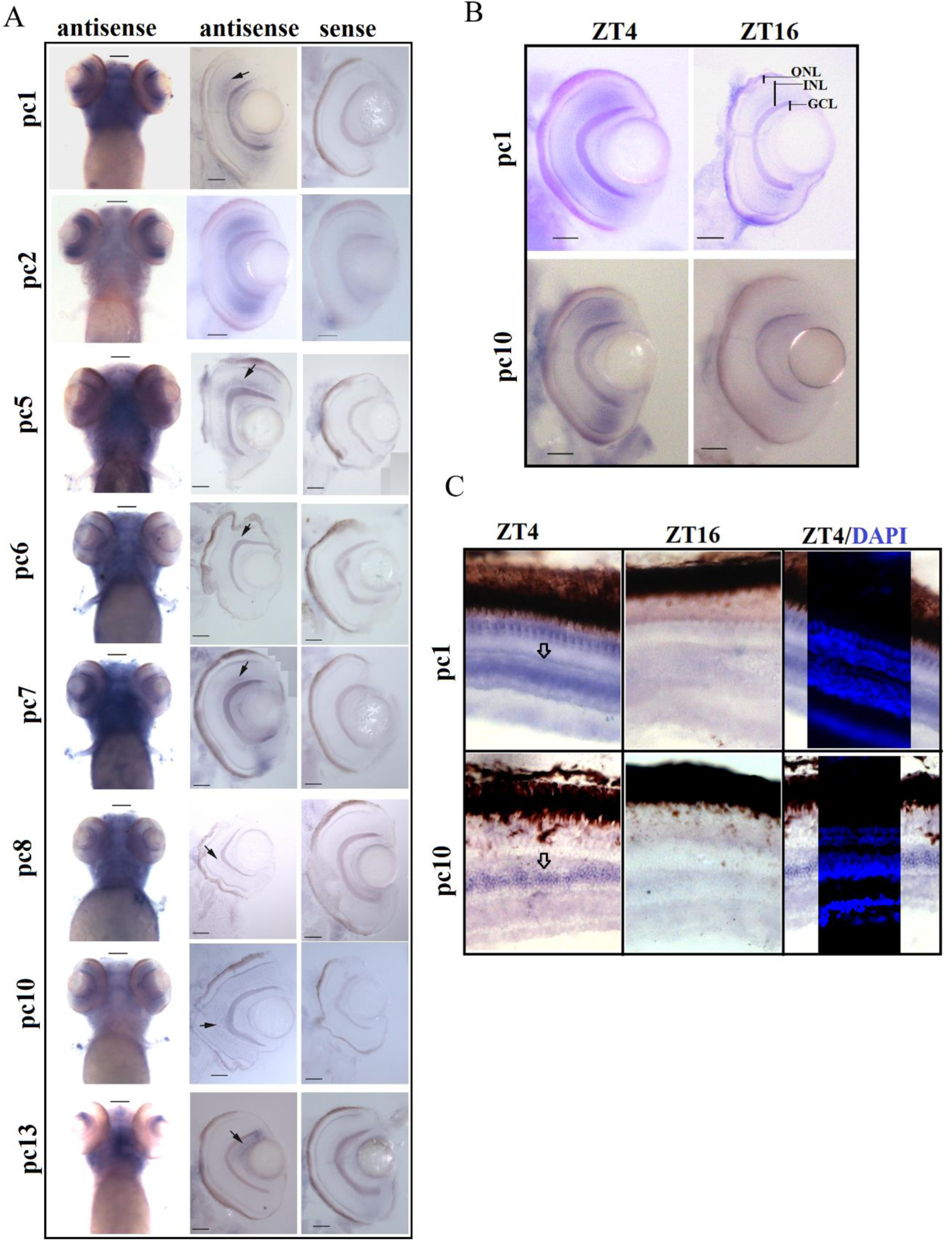
Novel transcripts are INL enriched revealed by RNA *in situ* hybridisation. **(A)** WISH on 5dpf zebrafish larvae shows eye-specific expression (left).WISH followed by cryosection (20μm thickness) revealed strong signal in INL; indicated by an arrowhead (middle), while sense probe shows no signal (right). ONL, outer nuclear layer (photoreceptor layer); INL, inner nuclear layer; GCL, retinal ganglion cell layer. (B) WISH on 5dpf larvae followed by cryosection indicate transcripts pc1 and pc10 differential expression between ZT4 and ZT16. (C) RNA *in situ* of pc1 and pc10 transcripts; on adult eye cryosection (20 μm thickness) collected at ZT4 (left), ZT16 (middle), ZT4 section counterstained DAPI reveal nuclear layers (right)

Spanjaard et al. recently reported the single cell expression profiling of zebrafish embryos at 5dpf (Spanjaard et al. 2018). We used this data to identify the cell type identity of differentially expressing transcripts identified in our study (Figure S2, S3). Although several novel transcripts were not detected due to the limited coverage of single cell transcriptome dataset, expression of pc2 and pc10 could be unambiguously assigned to retinal R1, R2 and R3 cluster of cells (Figure 6A, B). In agreement with our in situ hybridisation results, these cells are located in the INL of the retina. Further, pc10 expressing cells are also positive for Cabp5b, a marker of bipolar cells (Glasauer and Neuhauss 2016). In future, this expression overlap can be used for cell type-specific functional studies. Next, we used the single cell transcriptomics data to identify the cells of the retina where there is the highest convergence of expression of core clock genes. As shown in Figure 6C, the majority of the core clock genes show the highest expression in the R3 cell cluster, which also expresses pc2 and pc10.

**Figure 6:**
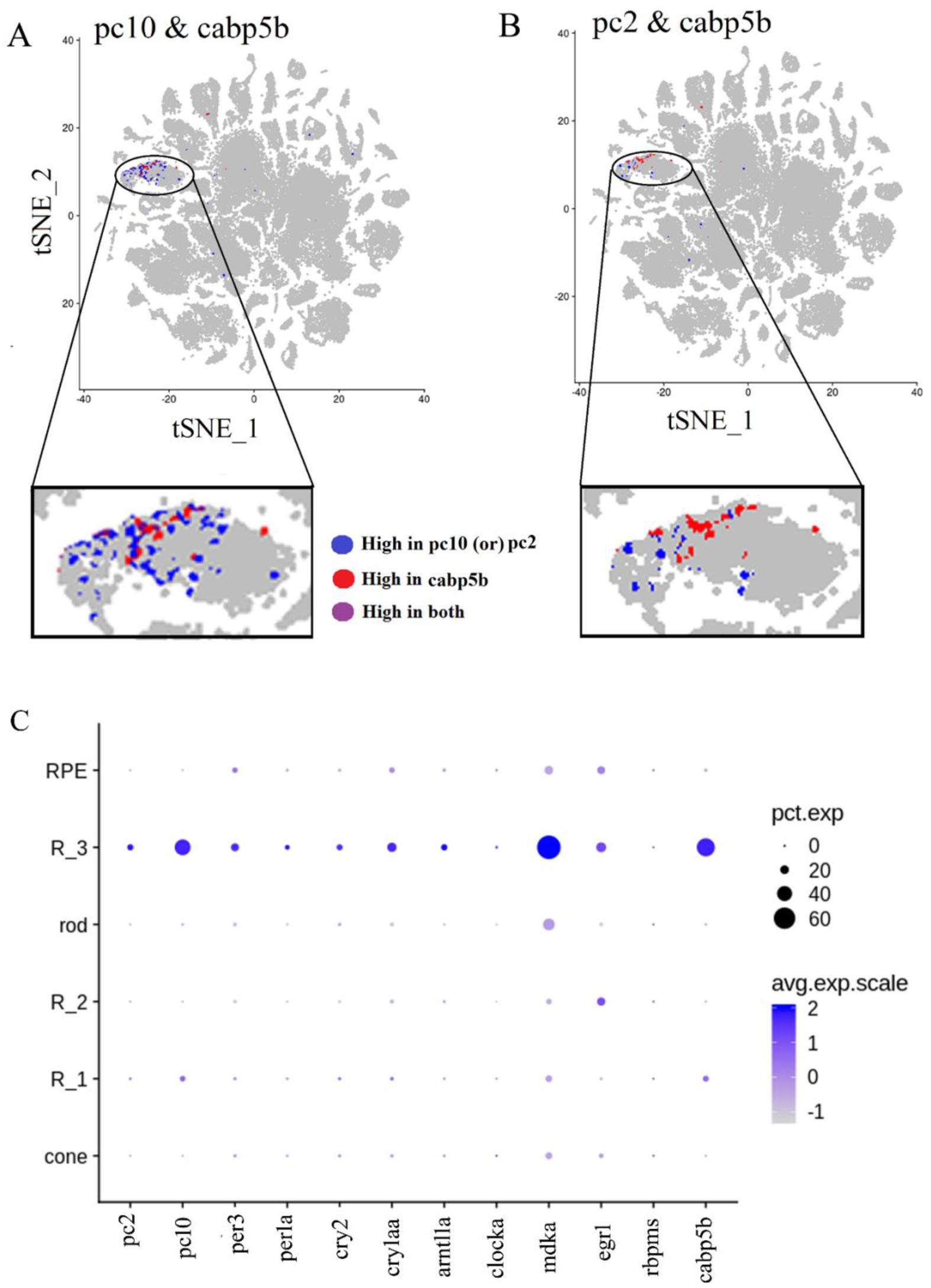
Cell type identity of novel transcripts using single cell transcriptome data: A, B) tSNE representation of pc1, pc2 and cabp5b expression in cell type clusters identified from 5dpf whole zebrafish single cell transcriptome. The enlarged version is to highlight the co-localization between novel transcripts and cabp5b, a bipolar cell marker. (C) Dotplot indicates the expression of novel transcripts (pc1 & pc10), core clock genes and marker genes in cell types clusters associated with retina (mdka – horizontal cells, erg1-amacrine cell, rbpms-retinal ganglion cells, cabp5b-bipolar cells), dot size indicates percentage of cells expressing gene of interest.

Endogenously produced dopamine and light are the two major factors known to regulate retinal diurnal and circadian physiology(Witkovsky 2004). Hence, we decided to understand the role of dopamine and light on the differential expression of novel transcripts. Dopamine-producing amacrine cells are present in the INL layer, the main source of dopamine in the retina, can be selectively ablated using a combination of 6-OHDA (6-hydroxydopamine) and Pargyline (Li and Dowling 2000). We used intra-ocular injections of a mixture of 6-OHDA and Pargyline (Figure 7B). We confirmed the ablation by immunohistochemical staining of Th1 (Figure 7C), a specific marker of dopaminergic amacrine cells. Bilateral uninjected control eye and the injected eye were compared for studying the effect of loss of dopaminergic neurons in the presence and absence of light. Fishes were deprived of light by extending the dark period for an additional four hours after the control fishes were transferred to light in the standard day-light cycle (Figure 7A). The positive control, per2 showed an evident up-regulation in the presence of light (Besharse et al. 2004). This regulation was abolished in the absence of amacrine cells but could be restored by external intra-ocular administration of dopamine, clearly showing the combined and mutually dependent effects of light and dopamine on per2 mRNA expression (Figure 7D). We then extended the analysis to all the 14 novel transcripts identified in our transcriptomics study and validated by qRT-PCR. None of the transcripts was affected by light or dopamine, suggesting that the circadian fluctuations seen in our experiments are driven by intrinsic regulation, independent of external cues (Figure 7E, F).

**Figure 7:**
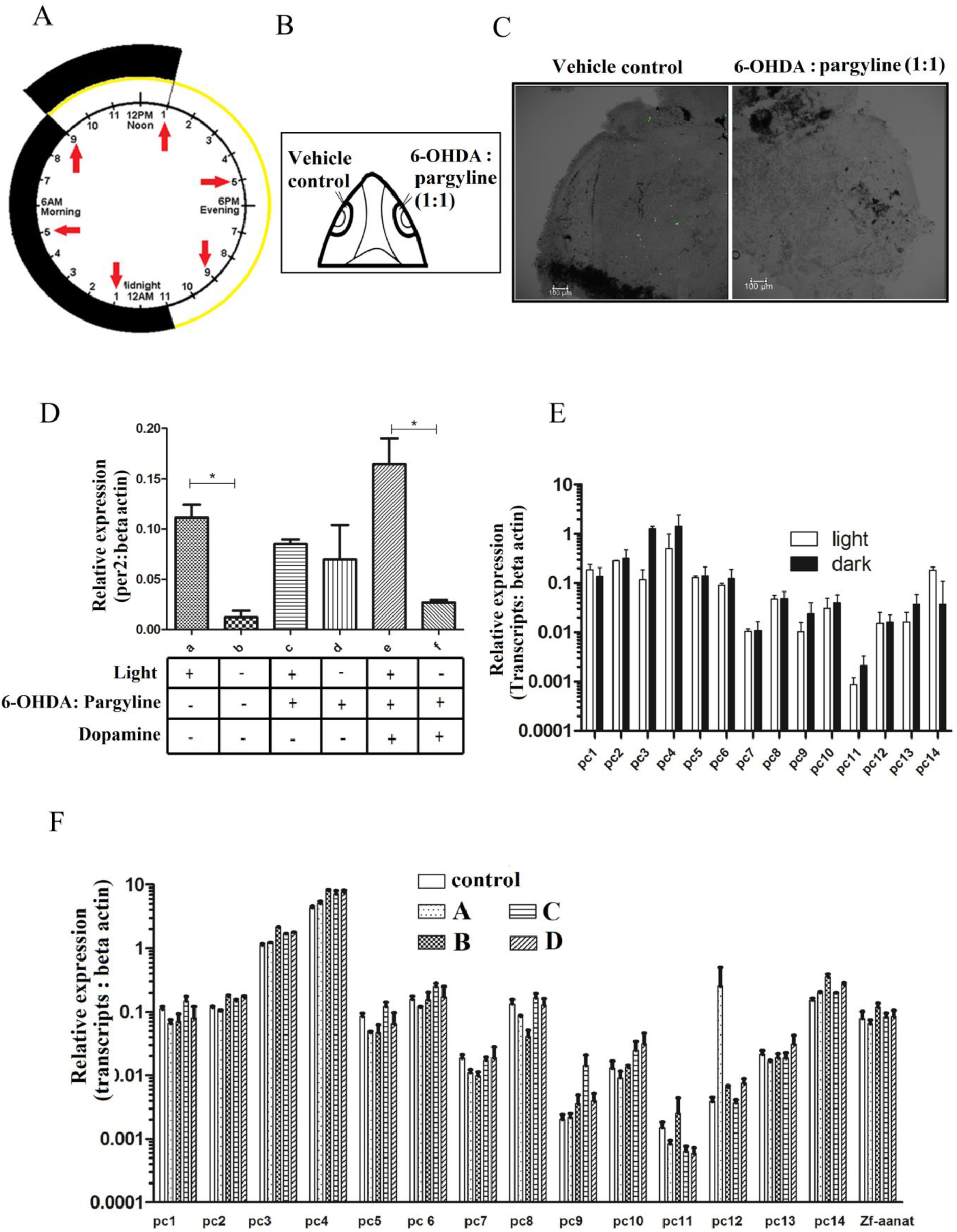
Expression of novel transcripts is unaffected by light and dopamine: (A)Schematic of ectopic light depletion regime;(B) Schematic of chemical ablation of Th+ amacrine cells by intra-ocular injection in adult zebrafish; (C) Th+ amacrine cells visualised in vehicle-injected and chemical ablated eye-cup by anti-Th+ immunolabeling on retinal whole mount preparation. (3D constructed Confocal micrograph); (D) zf-per2 relative expression from ZT4 harvested the whole eyecup along the following conditions; a (light), b (3hrs light depleted), c (light/Th+ amacrine cells ablated), d (3hrs light depleted /Th+ amacrine cells ablated), e (light/Th+ amacrine cells ablated/200μM dopamine injected at ZT0) and d (3hrs light depleted / Th+ amacrine cells ablated/200μM Dopamine injected at ZT0). The values, normalised for beta-actin, are mean ± s.e.m. (n=5); (E) Relative expression of novel transcripts from ZT4 measured between light and light deprived condition. The values, normalised for beta-actin, are mean ± s.e.m. (n=5); (F) Relative expression of novel transcripts from ZT4 measured between control (light/vehicle control), a (light/Th+ amacrine cells ablated), b (light/Th+ amacrine cells ablated/200μM dopamine injected at ZT0), c (light /Th+ amacrine cells ablated/200μM dopamine injected at ZT0) and d (3hrs light depleted / Th+ amacrine cells ablated/200μM Dopamine injected at ZT0) and. The values, normalised for beta-actin, are mean ± s.e.m. (n=5).

Next, we compared the expression pattern of these novel transcripts under constant darkness to verify that even without external light, the transcripts show rhythmic expression in the absence of light (Figure 8A). Total RNA isolated from whole eyecups from fishes maintained under constant dark conditions(Koike et al. 2012) (D/D) or entrained (L/D) condition were isolated and subjected to qRT-PCR analysis for known circadian genes (per2, aanat1 and aanat2) and novel transcripts identified in this study. Per2 was rhythmic only in response to light whereas aanat2 continues to show rhythmic expression even in constant darkness (Figure 8B). Amongst the novel transcripts seven (pc1, 3, 5, 6, 7, 8, 10 and11) peaked early in the day irrespective of the light cue while pc13 and 12 peaked during the early night. The latter two transcripts were arrhythmic under constant darkness (Figure 8C).

**Figure 8:**
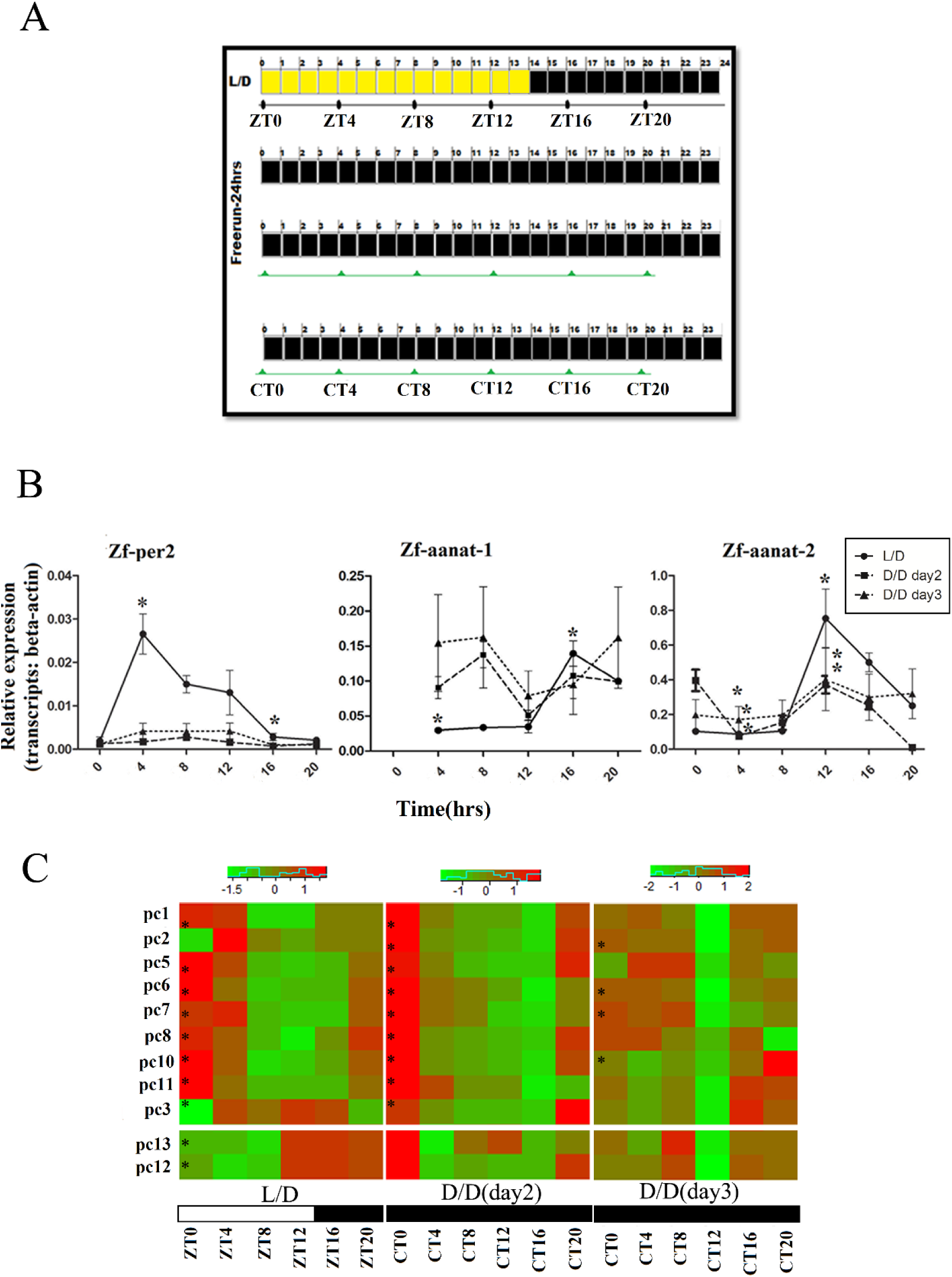
Circadian expression of novel transcripts: (A) Schematic showing the sampling time points between entrained (light/dark) and free run (dark/dark) conditions. (B) Relative expression of positive control genes in entrained and free run condition. The values, normalised for beta-actin, are mean ± s.e.m. (n=5). The X-axis denotes the time of collection in entrained (L/D) and free run condition (D/D) (C) Heat map represents the relative expression of novel transcripts in entrained (ZT) and free run conditions (CT). The values, normalised for beta-actin, are mean ± s.e.m. (n=5). * indicated the statistical significance of the transcripts rhythm calculated by meta2d.

## Discussion

The retina is an extension of the central nervous system, composed of a complex network of neurons and glia. By their very function, the cells of the retina undergo rapid, dynamic changes during the course of the day and night cycles. The ability of these cells to anticipate and respond rhythmically, yet tweaking the response according to seasonal changes in day-night cycles, is a critical factor for the survival of the organism. Unlike rodent models, the diurnal nature of zebrafish behaviour makes it an excellent model for human circadian biology. Further, our comparative assessment of core circadian gene peak expression between zebrafish and baboon placed zebrafish more closely to baboon.

Although the zebrafish retinal core clock component shows marked similarities to the mammalian retinal clock, it also differs in specific features. The genome duplication events in the evolutionary lineage of the zebrafish genome suggest that many circadian genes may have duplicated copies that are free to accumulate mutations and develop modified activity. This is even true for many zebrafish core circadian genes which exist as multiple paralogs (Liu et al. 2015). Surprisingly, our comparative assessment of the temporal expression of circadian genes in eye/retina of zebrafish, mouse and baboon revealed a previously unknown difference. Circadian genes in zebrafish(Huang et al. 2018) and mouse eye (Storch et al. 2007)show rhythmic expression, while all key circadian genes were arrhythmic in baboon retina (Mure et al. 2018) Similar non-circadian expression of Clock gene-a key circadian transcription factor was documented in human neurons and cortex(Fontenot et al. 2017; Chen et al. 2016). These differences hint towards an organ/organism-specific repurposing of core circadian genes whose biological significance is yet to be studied.

To the best of our knowledge, this is the first study to report the temporally resolved, circadian gene expression of adult zebrafish whole eye especially highlighting novel retinal transcripts. The sequencing of the human genome led to the surprising insight that only 2% of it codes for protein-coding genes while a significant, previously unknown fraction gives rise to transcripts that do not code for proteins. Currently, the core clock from insects and mammals has predominantly protein components, perhaps since the mechanistic studies on the circadian clock pre-dates the discovery of non-coding RNAs as a significant layer of gene regulation. It is intuitively appealing to speculate that the unbiased discovery of novel circadian transcripts, several of which do not have a strong coding potential may pave the way for a deeper understanding of the regulation of clock components. The novel transcripts reported here include two non-coding RNAs (pc1 and pc6), with absolutely no protein-coding potential or association with ribosomes in any tissue. In the light of recent evidence that many authentic long non-coding RNAs are associated with the ribosome (Carlevaro-Fita et al. 2016), it is premature to assume that all the ribosome-associated novel transcripts reported here give rise to proteins. Zebrafish is amenable to knockdown gene experiments to functionally characterise these transcripts in future. In future, these transcripts can be characterised for their role in circadian gene regulation and retinal physiology.

Besides these known genes, our analysis also revealed 18,000 apparently novel transcripts. It has been shown in several studies that the increased sequencing depth in NGS studies, can add a large number of fragmented, low expression transcripts that may be artifacts of the assembly process, and in any case are not amenable to further studies. Although the primary clock mechanism, limited to a handful of genes, is extensively studied, in recent years a series of transcriptomics studies have shown that an overwhelming number of genes are regulated by the circadian clock (Hughes et al. 2009). The novel players in circadian biology reported here may provide insights into the distinctive mechanisms in operation in the peripheral clocks. The retina is particularly interesting in this context since its input are necessary for SCN circadian rhythm (Paul et al. 2009).

Recent advancement in the single cell transcriptome rendered high confidence cell type catalogues on many organs including zebrafish retinal cells. Other than obtaining cell type identity for differentially expressed transcripts, our analysis also indicates the majority of the novel differentially expressed transcripts from zebrafish eyecups were expressed in the inner nuclear layer. Initially, it was thought that self-sustained circadian clocks of the photoreceptors in the retina drive rhythmic melatonin release. Ruan et al. showed that inner nuclear layer and ganglion cell layers express the core clock genes (Ruan et al. 2006) and in vitro cultured mouse retina continued to show sustained oscillations after several weeks in spite of the degradation of the photoreceptors. In agreement with the model proposed by Ruan et al., we find that majority of core clock genes in zebrafish retina were expressed in the inner nuclear layer based on the single cell transcriptome. Further, species-specific circadian transcripts may provide explanations for adaptations unique to the species and its habitat that tweak the basic clock mechanism to produce species-specific variations as in the case of blind cavefish (Cavallari et al. 2011).

In summary, we present the circadian transcriptome of zebrafish retina and identify several novel clock regulated genes. Spatially, many of these RNAs are expressed in the Inner Nuclear Layer, which has earlier been shown to harbour cell autonomous clocks in mice. The temporally resolved expression of these novel transcripts show that at least some of these transcripts show sustained oscillations even in the absence of external cues.

## Acknowledgement

The authors acknowledge the Council of Scientific and Industrial Research (CSIR), Government of India (HCP0008) for funding. The authors acknowledge Mr. Asgar Hussain Ansari for his critical inputs in RNA-seq analysis and R-packages used and Rijith Jayarajan for assistance in sequencing. We also acknowledge CSIR-IGIB zebrafish facility, imaging facility and high computing facility for the infrastructure provided.

## Conflict of interest

There are no conflicting interests to declare.

## Author contribution

S.R and B.P designed the experiments. S.K.V performed RNA-seq. B.R.I performed the RNA-seq analysis. S.R and S.S performed all zebrafish related experiments. S.R analyzed the single cell transcriptome dataset. B.P along with S.R and S.S were involved in data analysis, interpretation and writing of manuscript.

## Availability of data

The RNA-sequencing data have been deposited in Sequence Read Archive (SRA) under accession number SRP157045.

